# Seasonal variation in wind speed and oceanic salt spray favors delayed reproduction in coastal yellow monkeyflowers

**DOI:** 10.1101/2025.11.07.687229

**Authors:** Katherine Toll, Chloe Crockett, Lauren Tocci, David B. Lowry

**Affiliations:** University of South Carolina, Department of Biological Sciences, Columbia, SC; Michigan State University, Department of Plant Biology, East Lansing, MI; Plant Resilience Institute, Michigan State University, East Lansing, MI, 48824

**Keywords:** adaptation, coastal, wind, salt spray, flowering time, monkeyflower, Bodega Marine Reserve

## Abstract

**Premise:** The optimal timing of reproduction depends on the relative risk and benefit of continued growth. These risks and benefits depend on the local environment, which varies seasonally. While temperature and water availability are well studied selective agents on reproductive timing, less is known about other seasonally variable factors.

**Methods:** To test whether seasonal variation in wind and salt spray influences selection on reproductive timing, we performed an experimental manipulation of flowering time and ocean exposure in a coastal environment. Flowering plants from twelve yellow monkeyflower populations spanning a coastal-to-inland gradient were placed on a coastal bluff in spring and summer, which differ in average wind speed. A subset of plants were protected from wind and salt spray, and the remaining plants were exposed to these factors.

**Results:** In the windier spring cohort, ocean-exposed individuals had severe necrosis and produced few flowers and seeds, while protected plants had little-to-no necrosis, producing more flowers and seeds than exposed plants. In the calmer summer cohort, ocean-exposed individuals had less severe necrosis and produced comparable numbers of flowers as protected plants. However, only coastal populations produced appreciable numbers of seeds when exposed to the ocean, consistent with prior evidence that coastal ecotypes are locally adapted.

**Conclusions:** Overall, these results suggest that seasonal variation in wind and salt spray is a key selective factor contributing to population differences in flowering time.

## INTRODUCTION

The timing of reproduction is a key life-history transition that affects fitness. Life history theory predicts that the optimal timing of reproduction should depend on the costs of continued growth versus the benefits of growing for longer (Cohen 1976, Kozłowski 1992). Costs of continued growth include an increased risk of tissue loss or death prior to reproduction, while the benefits include an increase in size and offspring number. These relative risks and benefits depend on an organism’s environment, which is rarely constant. Seasonal variation in temperature, daylength, as well as water and resource availability can limit the growing season, and are major drivers of variation in the timing of life history transitions across environments and over time (Rathcke and Lacey 1985, Cleland et al. 2007). While temperature and water availability are best-studied, other local factors also determine an organism’s growing season and could generate divergent selection on reproductive timing across environments.

Organisms inhabiting coastal environments experience several unique abiotic stressors relative to inland environments, including exposure to wave action (Connell 1972), oceanic salt spray, sand abrasion, and sand burial (DiVittorio et al. 2020). These stressors limit growth and often result in pronounced patterns of coastal vegetation zonation (Oosting and Billings 1942). Salt spray is a dominant factor governing the distribution and abundance of plant species in coastal environments (Barbour and DeJong 1977, Du and Hesp 2020), though its direct role in the evolution of plant physiology, morphology, and phenology is less well understood (Ahmad and Wainwright 1976, Itoh 2021). Many plant lineages have evolved putative adaptations to coastal stressors, such as thick succulent leaves and prostrate growth (Turesson 1922, Boyce 1954, Lowry 2012), though similar phenotypes may be induced by salt spray (Wells and Shunk 1938, Boyce 1951, Griffiths 2006). Since new reproductive structures are produced at apical meristems, above vegetative tissues where wind and salt spray exposure is higher, flowers and fruits are often more susceptible to damage than leaves. Damage to reproductive meristems is costly when meristems are limited (Geber 1990), and this type of damage might favor the evolution of strategies to escape or tolerate recurrent injury (Wise and Abrahamson 2008).

Yellow monkeyflower (*Erythranthe guttata* [syn. *Mimulus guttatus*] species complex) populations grow across a coastal-to-inland gradient and differ in life-history, reproductive timing, and growth habit (Hall and Willis 2006, Lowry and Willis 2010). Coastal populations are typically perennials with prostrate growth habits that tend to flower in summer, whereas inland populations growing near ephemeral water sources are typically annuals with erect growth habits that tend to flower in spring, before the summer drought. In our study region, coastal Northern California, wind speeds are generally higher during the winter-spring rainy season compared to the relatively calmer summer and fall, and much higher near the coast (Gilliam 2002, García-Reyes and Largier 2012). Coastal populations are exposed to oceanic salt spray produced by wave action and distributed by wind (García-Reyes and Largier 2012, Du and Hesp 2020).

To test whether seasonal variation in wind and salt spray might be a selective agent contributing to differences in life history timing between coastal and inland populations, we experimentally manipulated flowering time and ocean exposure in the field. We induced flowering using supplemental light, then placed plants (just prior to or after anthesis) outdoors at two timepoints, aligned with when inland annual (spring) and coastal perennial (summer) plants flower at the same latitude. We protected a subset of plants from salt spray and wind in both seasons with exclosures and exposed the remaining plants in control structures. If coastal populations are able to escape tissue death and increase fitness by temporally escaping wind and salt spray, we predicted that (1) tissue death would be higher and fecundity would be lower during the windier spring timepoint compared to calmer summer timepoint, (2) protection from wind and salt spray would reduce tissue death and increase fitness, and (3) the benefit of protection would be greater in the spring than in the summer.

## MATERIALS AND METHODS

We performed our field experiment at the Bodega Marine Reserve in Bodega Bay, Sonoma County, California, USA (38.31841, −123.07329). To prevent gene flow between experimental and local plants, we placed our arrays near, but not in, natural seeps containing monkeyflowers. Our site was 0.24 km and 0.47 km from two natural seep populations, and was within a comparable distance to the ocean as these natural populations (under 50 m).

We grew a single outbred family from each of 12 populations (Figure 1A, Appendix S1: Table S1). Outbred families were generated by intercrossing two different individuals from each population. Four populations were inland annuals from Sonoma County, California, and four populations were coastal perennials spanning a latitudinal cline in plant height from Oregon to Sonoma County (Zambiasi and Lowry 2024). We also grew perennial populations from localities not directly exposed to the ocean: two near-coastal perennials from Sonoma County, and an inland perennial from a large river in Siskiyou County, California. Lastly, we included a near-coastal annual population that was collected under 1.6 km from the near-coastal perennials.

**Figure 1.**
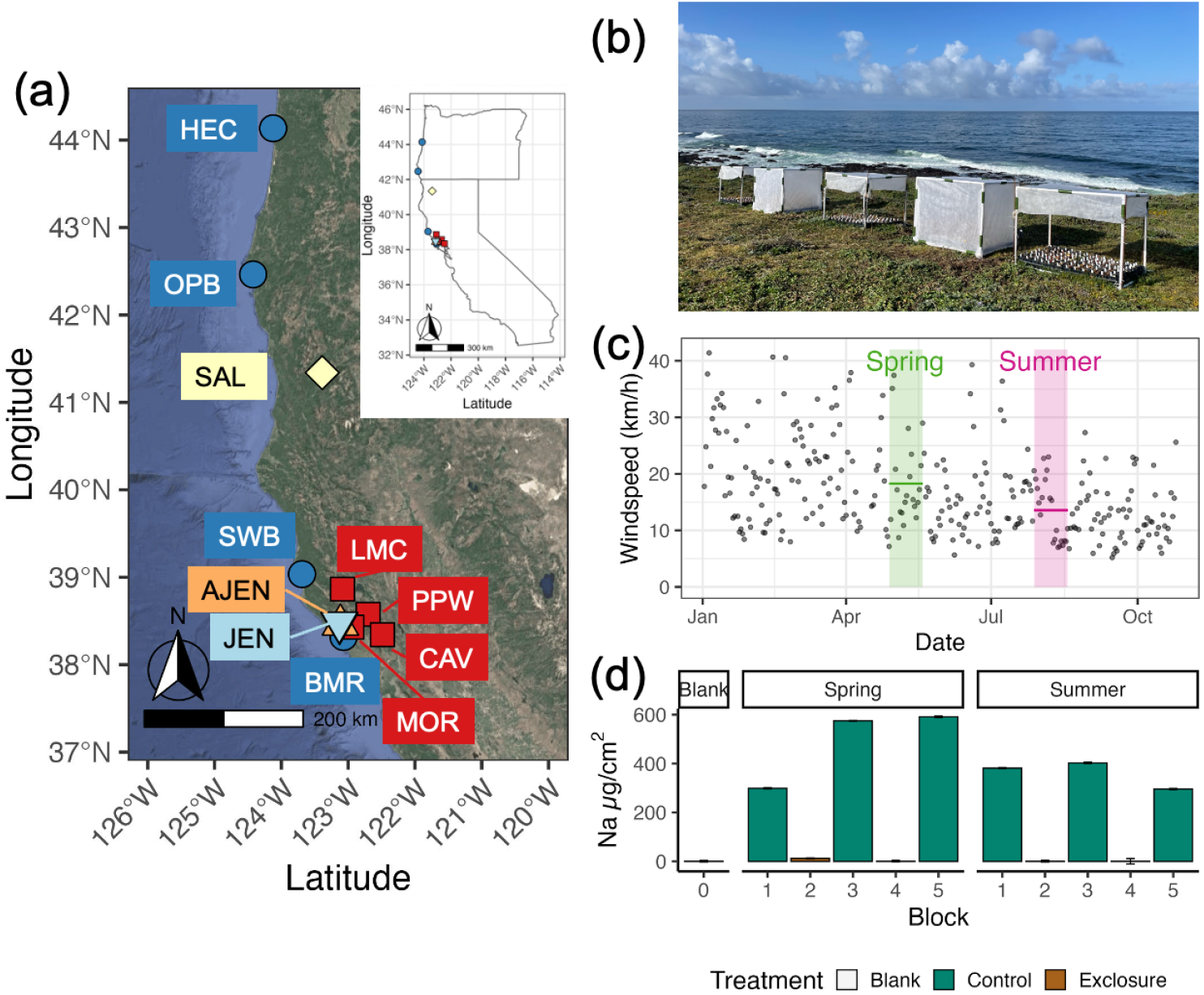
(a) Map of collection localities of coastal perennial (dark blue circles), inland annual (red squares), inland perennial (yellow diamond), near-coastal annual (orange triangle), and near-coastal perennial (light blue triangles) populations used in the experiment, (b) photograph of experiment at Bodega Marine Reserve (photo credit: K Toll), (c) plot of average daily wind speed at the Bodega Marine Reserve in 2023, and (d) plot of sodium accumulation on filter paper traps during the spring and summer exposure periods in exclosures (brown) and controls (green). Wind speeds were higher in the spring cohort compared to the summer cohort (c, pink vs green), and exclosures significantly reduced Na accumulation in both seasons (d, green vs brown).

We planted perennial and annual seeds a week apart to synchronize their development, following Popovic and Lowry (2020), in the Bodega Marine Reserve greenhouse (Spring perennials: 3/2/2023, annuals: 3/9/2023; Summer perennials: 6/5/2023, annuals 6/12/2023). We evenly distributed 9 seedlings per population across five blocks of six flats, for a total of 45 seedlings per population per cohort (540 plants per cohort). In each randomized block, six flats contained at least one seedling per population, while two flats were randomly assigned a second seedling. Since spring day lengths at our study site were not long enough to induce flowering (Friedman and Willis 2013), we grew plants under supplemental lights for 16 hours both seasons.

To protect plants from wind and salt spray in each cohort, we completely covered two out of five blocks with agrofabric (Figure 1B). Agrofabric exclosures were 108 cm x 84 cm x 87 cm and constructed of 3/4” PVC pipe covered with medium-weight agrofabric (Pro-34 1.0 oz/yd^2^ with 70% light transmission; OBC Northwest, Inc., Canby, OR, USA) affixed with hot glue and PVC clips. Each structure was secured to the ground with rebar attached to the PVC with zip-ties. Exclosures had one door facing away from the ocean secured with PVC clips to allow access to the plots. To control for agrofabric shading or moisture collection, the remaining three blocks were constructed identically, except that agrofabric only covered their tops and 30 cm down each side. In both seasons, we covered the control structures with bird netting to reduce herbivory. We did not observe herbivory in spring, but in summer, the inflorescences of a few plants on the edges of the control structures were eaten by mule deer (*Odocoileus hemionus*).

At the start of each cohort, we measured plant height, removed petals from any open flowers, and then moved flats into their respective outdoor blocks. Plants were left outdoors for three weeks in each cohort (spring: 4/28/2023 to 5/19/2023, summer: 7/28/2023 to 8/18/2023). We censused plants daily and manually self-fertilized up to three new flowers for each plant. Thereafter, we removed petals from each new open flower daily to prevent gene flow with local populations. We recorded leaf and inflorescence necrosis (tissue death) on a categorical scale weekly. Since we observed apical necrosis on ocean-exposed plants, we measured living tissue height on each plant weekly. We bottom watered flats as needed throughout each cohort. At the end of each cohort, we transported flats back into the greenhouse, where we collected hand-pollinated fruit individually as they ripened. Seeds from each fruit were counted by eye.

To quantify wind speed in each season, we downloaded average daily wind speed data from the Bodega Ocean Observing Node, measured within meters of our experiment (https://boon.ucdavis.edu/data-access/products/met/bml_wind_speed_daily). To quantify salt spray, we constructed filter paper traps with a 3D printer. The traps were 60 x 60 x 103 mm and had a 20 x 30 mm opening for filter paper on all four sides. We cut Whatman Filter paper (qualitative, grade 2) in 38 x 60 mm pieces and placed them on three sides of each trap. We affixed the traps to the southeast corner of each structure, 380 mm from the ground, and oriented them so one side faced west at the Pacific Ocean. West-facing filter paper samples were cut into a 1cm^2^ square using a pair of clean scissors that were washed between samples. The filter paper samples were then eluted in 10 ml of ultrapure 18MOhm water for 72 hours. An aliquot of that solution was taken out and spiked with Indium (internal standard to correct for drift). The samples were run on a THERMO ELEMENT2 HR-ICPMS at high-resolution mode. Sodium concentrations were calculated with external calibration of Na pure solution, spiked with In.

### Statistical Analysis

All statistical analyses were performed in R version 4.4.1 (R Core Team 2024). Mixed models were analyzed using *glmmTMB* (Brooks et al. 2017) *or ordinal* (Christensen 2023), model term significance was assessed using analysis of deviance (Wald *χ2* test) with *car* (Fox and Weisberg 2018), and pairwise comparisons were conducted using *emmeans* (Lenth et al. *2020)*. We predicted means and 95% confidence intervals using *ggeffects* (Lüdecke 2018).

To test whether necrosis severity differed between ocean-exposure treatments and season, we analyzed two necrosis proxies. Our first necrosis proxy was living tissue height over time, since we observed that spring ocean-exposed plants initially exhibited necrosis at their apical meristems and died back towards their bases. We first visualized height over time by plotting raw height values during each census. To reduce model complexity, we then calculated the change in living tissue height for each plant by subtracting the initial height measured in the greenhouse from the final height 21 days after being outdoors. We analyzed the change in height using a linear mixed model with population *(n* = 12), season (*n* = 2), and exclosure treatment (*n* = 2), all two-way interactions, and the three-way interaction as fixed effects, and flat (*n* = 60) nested within block (*n* = 10) as random effects. Our second necrosis proxy was a qualitative assessment of necrosis to leaves and inflorescences in three categories of severity (no, partial, or complete necrosis). We plotted the categories of severity over time, and tested whether leaf or inflorescence necrosis at the end of each exposure period differed between seasons and treatments using a cumulative link mixed model. The necrosis severity model consisted of season and exclosure treatment as fixed effects, and block as a random effect since interactive models and models with flat nested within block were singular.

To test whether fecundity differed between ocean-exposure treatments and season, we analyzed flower production and seeds produced by up to three hand-pollinated flowers per plant. We analyzed flowers per plant using a mixed model with a negative binomial error distribution since the poisson model was overdispersed. The model consisted of season, population, exclosure treatment, all two-way interactions, and the three-way interaction as fixed effects, and flat nested within block as random effects. To analyze seed production, we first averaged seeds produced by each plant to avoid pseudoreplication. We analyzed seeds per flower using a mixed model with a zero-inflated negative binomial error distribution since the Poisson model was significantly overdispersed. Because the model with a three-way interaction did not converge, the seed model consisted of season, population, exclosure treatment, and all two-way interactions as fixed effects, and flat nested within block as random effects.

## RESULTS

Average daily wind speeds were five km/h faster during the spring cohort compared to the summer cohort (spring mean (range) = 18.3 (7.2-37.4) km/h; summer mean (range) = 13.6 (7.0-23.0) km/h; Figure 2a). Sodium (Na) deposition in the three control structures ranged from 298.9-591.3 µg/cm^2^ in spring and 295.6-402.8 µg/cm^2^ in summer (Figure 2b). In each season, the two exclosures had little to no Na deposition (12.0 and 0.6 µg/cm^2^ in spring; 0.2 and 0.4 µg/cm^2^ in summer), comparable to a blank filter paper control (0.2 µg/cm^2^). Agrofabric exclosures effectively reduced Na accumulation (Figure 1d, 242 vs 3.3 µg/cm^2^, two-tailed *t*-test: *t* = 7.89, *df* = 5.03, *p*-value < 0.001). An intense spring storm briefly detached the bottom northwest corner of agrofabric covering one of the exclosures, reflected by a slightly higher Na concentration (12.0 µg/cm^2^) in that block, which is still 25 times lower than the control blocks. We promptly secured the agrofabric with additional adhesive and PVC clips after the storm.

**Figure 2.**
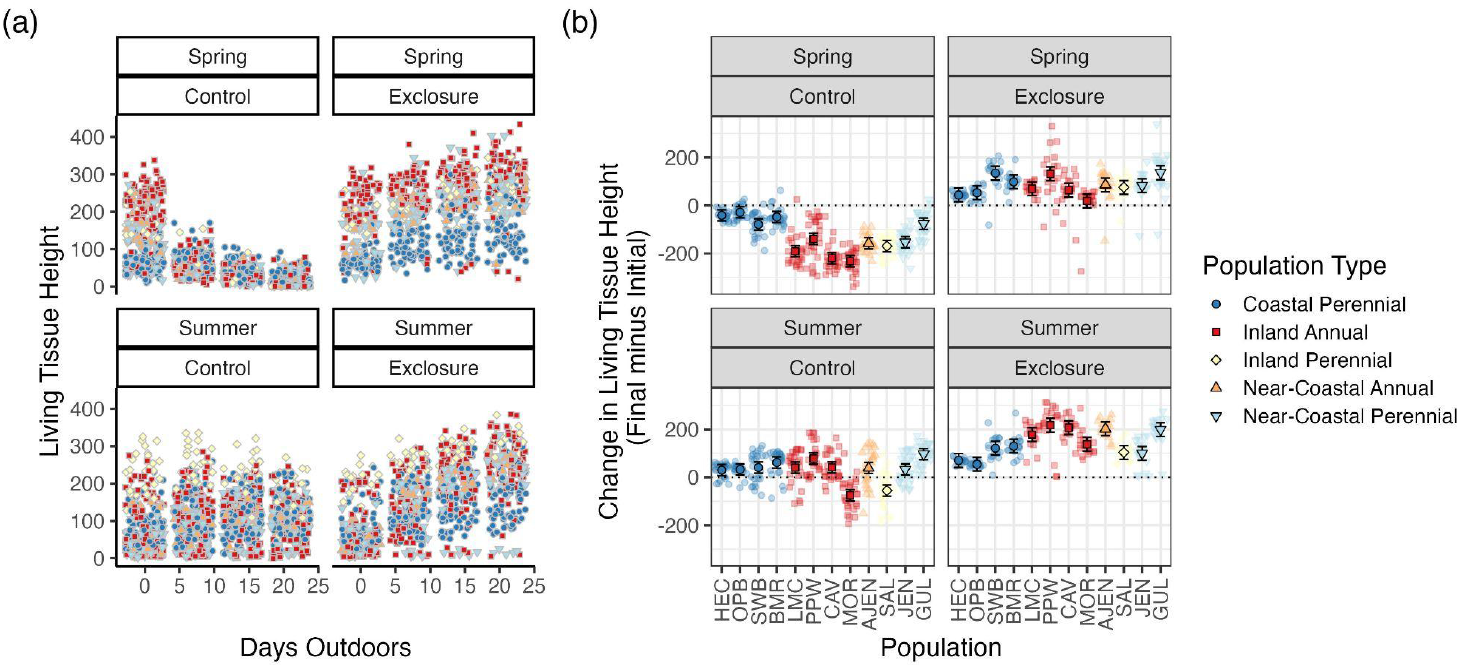
Spring flowering plants experienced severe apical tissue death when exposed to the ocean, reflected by the decrease in living tissue height, and agrofabric exclosures protected plants from necrosis. (a) Living tissue height over time, measured prior to moving plants outside and at three weekly censuses in ocean-exposed controls or agrofabric protected exclosures. (b) Individual height differences (final minus initial height) (background points) and predictions from a linear mixed model (black outlined points; mean and 95% confidence intervals).

Necrosis, estimated by living tissue height, was significantly associated with population, season, exclosure treatment, a population by season interaction, a population by treatment interaction, and the three-way population by season by exclosure treatment interaction (all *p* < 0.001, Table 1). In spring, when wind speeds were highest (Figure 1c), ocean-exposed plants experienced severe apical tissue death, reflected by a significant decrease in living tissue height in the control structures (Figure 2). In summer, we did not observe extreme apical necrosis and most ocean-exposed plants increased in height over time (Figure 2), but not as much as in the exclosures (all *p* ≤ 0.04, Appendix S1: Table S2). In both seasons, agrofabric-protected plants increased in height (Figure 2), and the change in height significantly differed between ocean-exposed and protected treatments (all *p* < 0.001, Appendix S1: Table S2).

**Table 1.** Analysis of Deviance Tables (Type III Wald χ2 tests) for change in height, leaf necrosis, inflorescence necrosis, flower production and seed production.

| Response | Factor | $\chi^2$ | df | p |
| --- | --- | --- | --- | --- |
| Change in height | (Intercept) | 11.9221 | 1 | 0.0005547 |
|  | Population | 545.0121 | 11 | < 2.2E-16 |
|  | Season | 17.7024 | 1 | 2.58E-05 |
|  | Treatment | 20.3557 | 1 | 6.43E-06 |
|  | Population:Season | 219.1006 | 11 | < 2.2E-16 |
|  | Population:Treatment | 206.2958 | 11 | < 2.2E-16 |
|  | Season:Treatment | 2.8391 | 1 | 0.0919932 |
|  | Population:Season:Treatment | 33.961 | 11 | 0.000367 |
| Leaf Necrosis | Season | 73.622 | 1 | < 2.2E-16 |
|  | Treatment | 129.923 | 1 | < 2.2E-16 |
| Inflorescence Necrosis | Season | 204.87 | 1 | < 2.2E-16 |
|  | Treatment | 362.99 | 1 | < 2.2E-16 |
| Flower production | (Intercept) | 11.788 | 1 | 0.0005962 |
|  | Season | 44.699 | 1 | 2.30E-11 |
|  | Population | 92.857 | 11 | 4.58E-15 |
|  | Treatment | 54.457 | 1 | 1.59E-13 |
|  | Season:Population | 30.849 | 11 | 0.0011638 |
|  | Season:Treatment | 25.86 | 1 | 3.67E-07 |
|  | Population:Treatment | 104.645 | 11 | < 2.2E-16 |
|  | Season:Population:Treatment | 35.268 | 11 | 0.0002238 |
| Seeds per flower | (Intercept) | 149.612 | 1 | < 2.2E-16 |
|  | Season | 120.804 | 1 | < 2.2E-16 |
|  | Treatment | 154.401 | 1 | < 2.2E-16 |
|  | Population | 361.07 | 11 | < 2.2E-16 |
|  | Season:Treatment | 133.033 | 1 | < 2.2E-16 |
|  | Season:Population | 25.824 | 11 | 0.006892 |
|  | Treatment:Population | 300.598 | 11 | < 2.2E-16 |

Leaf and inflorescence necrosis were both significantly associated with season (*p* < 0.001) and exclosure treatment (*p* < 0.001, Table 1). By the end of the spring cohort, 97-99% of ocean-exposed plants had complete leaf (314/324) and inflorescence (323/324) necrosis, i.e. no living tissue remaining, while the remainder had partial necrosis (some living tissue) (Appendix S1: Figures S1, S2a & S3, Table S3). By the end of the summer cohort, 78-83% of ocean-exposed plants had partial leaf (268/324) and inflorescence (253/324) necrosis, while the remainder had no necrosis (Appendix S1: S2b). A small percentage (9-12%) of plants, located near the northwest corners of the spring exclosures had leaf (26/216) and inflorescence (20/216) necrosis due to an intense storm that briefly detached the protective agrofabric.

Flower production was significantly associated with season, population, exclosure treatment, and all two-way and the three-way interactions (all *p* < 0.001, Table 1; Figure 3a). In spring, all ocean-exposed populations produced significantly fewer flowers than those protected in agrofabric exclosures (4-14 fewer flowers, all *p* < 0.001; Appendix S1: Table S4, Figure S4a & S5). In summer, some (7/12) ocean-exposed populations produced significantly fewer flowers than those protected in agrofabric exclosures, and the difference was smaller (2-7 fewer flowers except one inland annual that made 24 fewer flowers, *p* ≤ 0.05; Appendix S1: Table S4, Figure S4a). Ocean-exposed populations produced 2-7 fewer flowers in spring compared to summer (*p* < 0.001, Appendix S1: Table S4, Figure S4a). Most (8/12) agrofabric protected populations did not differ in flower number between seasons, and significant differences varied in direction (−14 to +8 in spring vs summer, Appendix S1: Table S4, Figure S4a).

**Figure 3.**
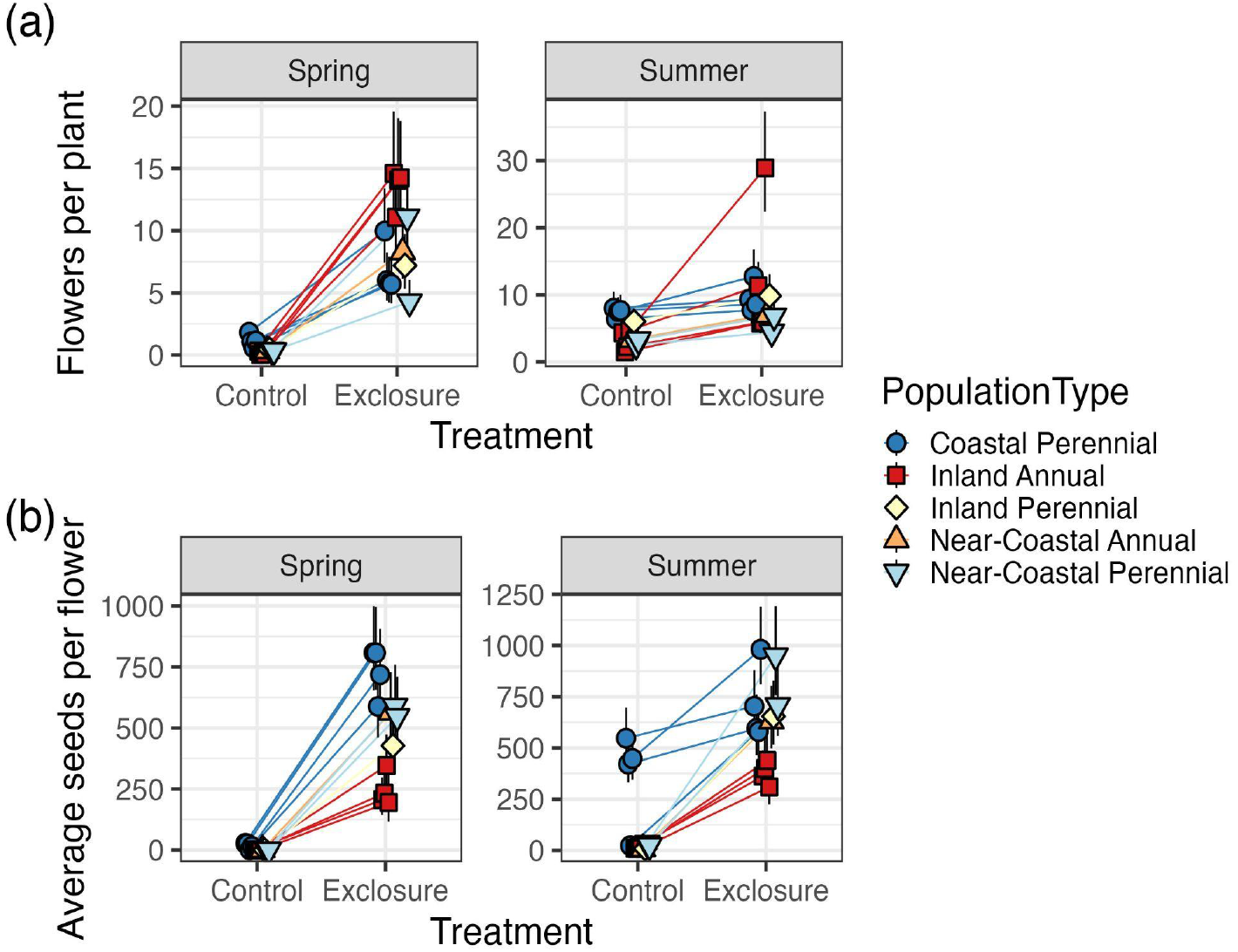
Plants produced fewer flowers per plant (a) and seeds per flower (b) in spring compared to summer when exposed to the ocean, plants protected by agrofabric exclosures produced more flowers and seeds than those exposed to the ocean, and the benefit protection was greater in spring. (a) Predicted flowers per plant (mean and 95% confidence intervals), and (b) predicted seeds per flower (mean and 95% confidence intervals) from mixed models.

Average seed production per hand-pollinated flower was significantly associated with season, population, exclosure treatment, and all two-way interactions (all *p* ≤ 0.006, Table 1; Figure 3b). In spring, only coastal perennials produced seeds when exposed to the ocean (Appendix S1: Figure S4). Spring ocean-exposed populations produced 192-772 fewer seeds per flower than those protected in agrofabric exclosures (all *p* < 0.001, Appendix S1: Table S5, Figure S4b). In summer, most (10/12) ocean-exposed populations produced fewer seeds compared to those protected in agrofabric exclosures (297-902 fewer seeds per flower, *p* < 0.001, Appendix S1: Table S5, Figure S4b). Most ocean-exposed plants that produced seeds in summer were coastal perennials (Appendix S1: Figure S6). Ocean-exposed annuals and the inland perennial population did not differ in seeds per flower between seasons, largely because they produced almost no seeds (*p* ≥ 0.1, Appendix S1: Table S5, Figure S4b). In contrast, coastal and near-coastal perennials exposed to the ocean produced 22-513 fewer seeds per flower in spring compared to summer (*p* ≤ 0.03, Appendix S1: Table S5, Figure S4b). Almost all (11/12) agrofabric-protected populations did not significantly differ between seasons in seeds per flower (*p* ≥ 0.09, except one near-coastal perennial *p* = 0.03, Appendix S1: Table S5, Figure S4b).

## DISCUSSION

The timing of flowering is a key life-history transition that affects plant fitness. The optimal timing of reproduction depends on the costs and benefits of continued growth versus reproduction. These costs and benefits vary depending on an organism’s environment, which often varies seasonally. Although much is known about the role of temperature and precipitation in driving variation in phenology, less is known about the role of hyper-local factors like wind and oceanic salt spray. Inland annual yellow monkeyflower populations flower in early spring to avoid the summer drought, whereas coastal perennial populations inhabit seeps where moisture is available year-round and flower in the summer (Hall and Willis 2006, Lowry and Willis 2010). Experimentally inducing spring flowering typical of inland populations at a coastal site resulted in high rates of necrosis, low flower production and low seed set in all populations. In contrast, experimentally inducing late summer flowering typical of coastal populations at the same coastal site resulted in lower rates of necrosis, higher flower production, and higher seed set compared to spring. Almost all individuals that produced seed when exposed to the ocean were coastal perennials, consistent with prior studies demonstrating that these populations are locally adapted (Hall and Willis 2006, Lowry et al. 2008, Popovic and Lowry 2020). In both seasons, individuals protected from wind and salt spray had almost no necrosis (Figure 2), and produced more flowers and seeds than ocean-exposed individuals (Figure 3). The benefit of protection was greater in spring than in summer (Figure S2). Collectively, our results suggest that escape from tissue death is a major benefit of summer flowering in coastal populations.

Plants inhabiting coastal environments experience multiple above-ground stressors, including wind and resultant salt spray and sand abrasion (Du and Hesp 2020). Above-ground meristems that produce new leaves or flowers and fruits are especially sensitive to these abiotic stressors (Boyce 1954). When meristems are limiting, selection should favor strategies to either escape or tolerate recurrent meristem damage (Geber 1990, Wise and Abrahamson 2008).

Coastal perennial monkeyflowers likely use multiple strategies to protect their meristems from wind and salt spray, including keeping their meristems low to the ground via prostrate growth; salt spray intensity decreases with proximity to the ground (Martin 1959, Barbour 1978, Griffiths 2006, Zambiasi and Lowry 2024). Yet these strategies were insufficient to rescue spring fecundity in the populations we examined, suggesting that ocean-exposed populations are constrained to reproducing during relatively still periods of the year.

In our study region, northern coastal California, wind speeds vary seasonally and are generally highest during upwelling (April-June), weakest from July to September, and highly variable during frequent storms from December through February (García-Reyes and Largier 2012). Heavy winter precipitation causes rapid leaching of salt to deeper layers of sand, ameliorating the toxic effects of soil salinity (Du and Hesp 2020). While our study focused on temporal variation at an ocean-exposed shoreline, perennial monkeyflowers also inhabit areas near the coast that are protected from the ocean. Protected populations can flower during the windy spring season (Toll, pers. obs.), potentially because their reproductive structures are not recurrently damaged. A common garden study found that ocean-protected perennial populations are taller than nearby ocean-exposed populations, but did not find differences in flowering time, suggesting that field phenology differences may be due to plasticity (Zambiasi and Lowry 2024). Nevertheless, height divergence may be facilitated by assortative mating within protected and exposed forms (Aston and Bradshaw 1966, Daehler et al. 1999, Foster et al. 2007, James et al. 2023, Itoh et al. 2024).

Removing wind and salt spray with exclosures dramatically decreased necrosis and increased fecundity. We were unable to directly separate these two factors, since high winds are ultimately the cause of salt deposition. These factors likely operate interactively, since salt spray penetration increases when plant tissues are abraded by wind-blown sand (Du and Hesp 2020). However, we have only observed apical necrosis in natural populations of coastal monkeyflowers directly exposed to the ocean and not in nearby protected populations (Appendix S1: Figure S7), consistent with Wells and Shunk’s (1938) observations of shoot injury to woody plants near the ocean but not in nearby freshwater estuaries where wind speeds are similar. Additionally, we have only observed aerial necrosis at ocean-exposed sites in multiple reciprocal transplant experiments with yellow monkeyflowers (Lowry et al. 2008, Toll et al. 2024). Although the relative importance of wind versus salt spray in structuring plant communities was debated (Wells and Shunk 1938, Oosting and Billings 1942, Boyce 1954), the general consensus is that both factors are important in determining the distribution, abundance, and phenotypes of organisms in coastal environments. Our study adds to a growing number of recent studies directly testing the ecological significance of traits underlying adaptation to coastal environments (Wilkinson et al. 2021, Itoh et al. 2024).

## ACKNOWLEDGEMENTS

We would like to thank Jackie Sones and Luis Morales for facilitating research at the Bodega Marine Reserve. Wind speed data was provided by the University of California, Davis, Bodega Marine Laboratory. Veronica Toll assisted with field data collection. Madi Stessman designed and 3D printed the filter paper traps for salt spray quantification. Micheal Bizimis and Jim Sexton from the University of South Carolina Center for Elemental Mass Spectrometry quantified sodium samples. C.C. and L.T. were supported by the University of South Carolina Honors College Research Grant. This work was funded by a National Science Foundation Division of Integrative Organismal Systems Grant to D.B.L. (IOS-2153100).

## AUTHOR CONTRIBUTIONS

Katherine Toll: Methodology, Investigation, Formal Analysis, Data Curation, Writing - Original Draft, Writing - Review & Editing, Visualization. Chloe Crockett: Investigation. Lauren Tocci: Investigation. David B. Lowry: Conceptualization, Methodology, Writing - Review & Editing, Resources, Supervision, Funding acquisition.

## Notes

### Competing Interest Statement

The authors have declared no competing interest.

