## Appendix S1 for "Seasonal variation in wind speed and oceanic salt spray favors delayed reproduction in coastal yellow monkeyflowers"

**Toll et al. – Appendix S1**

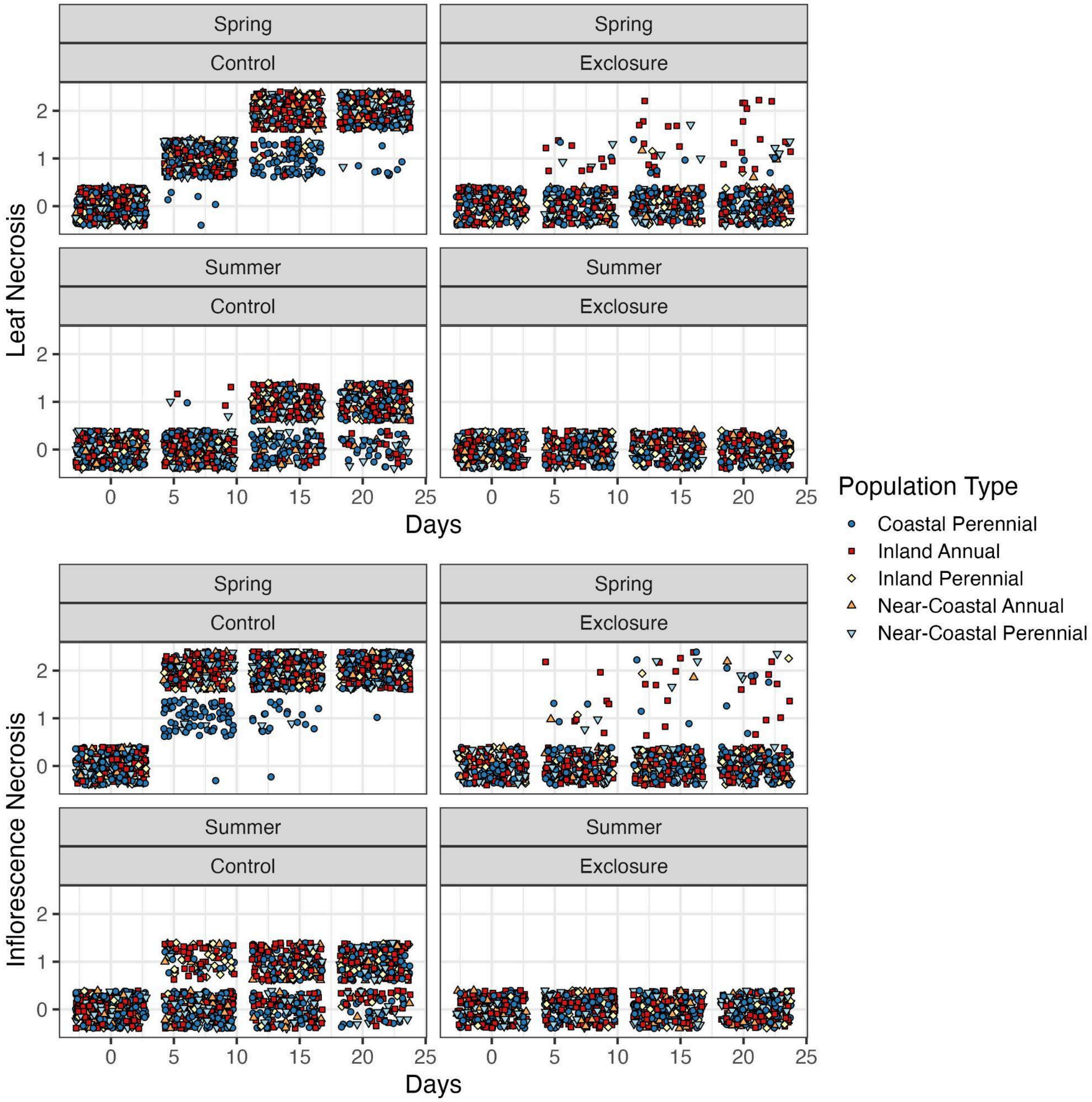


**Figure S1**. Plants in the spring cohort had more severe necrosis, and agrofabric exclosures effectively protected plants from necrosis. Necrosis was assessed in three categories of severity (no [0], partial [1], complete [2]). Necrosis was assessed prior to moving plants outside from the greenhouse at the Bodega Marine Reserve (Spring: 4/28/23; Summer: 7/28/23), and during three weekly censuses while plants were outdoors in control (ocean-exposed) structures or agrofabric-covered exclosures (Spring: 5/5/23, 5/12/23, 5/19/23; Summer: 8/4/23, 8/11/23, 8/18/23).

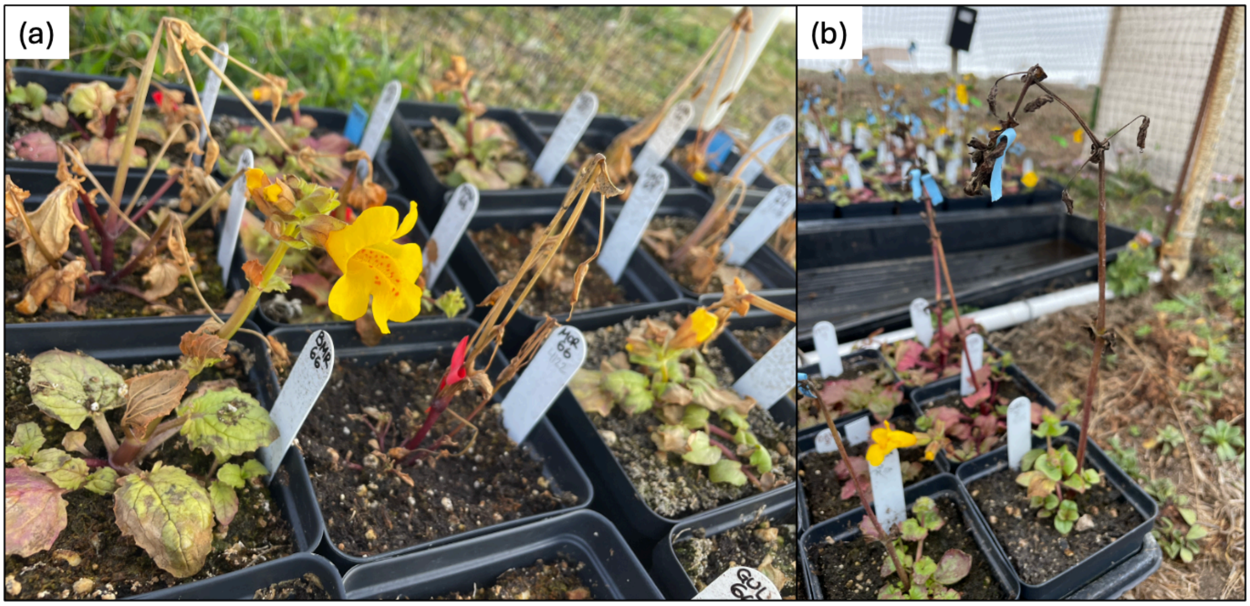


**Figure S2.** Photographs of necrosis in (a) spring plants one week after ocean exposure and (b) summer plants three weeks after exposure. (a) depicts a local flowering coastal perennial accession in the foreground next to severely necrotic annuals.

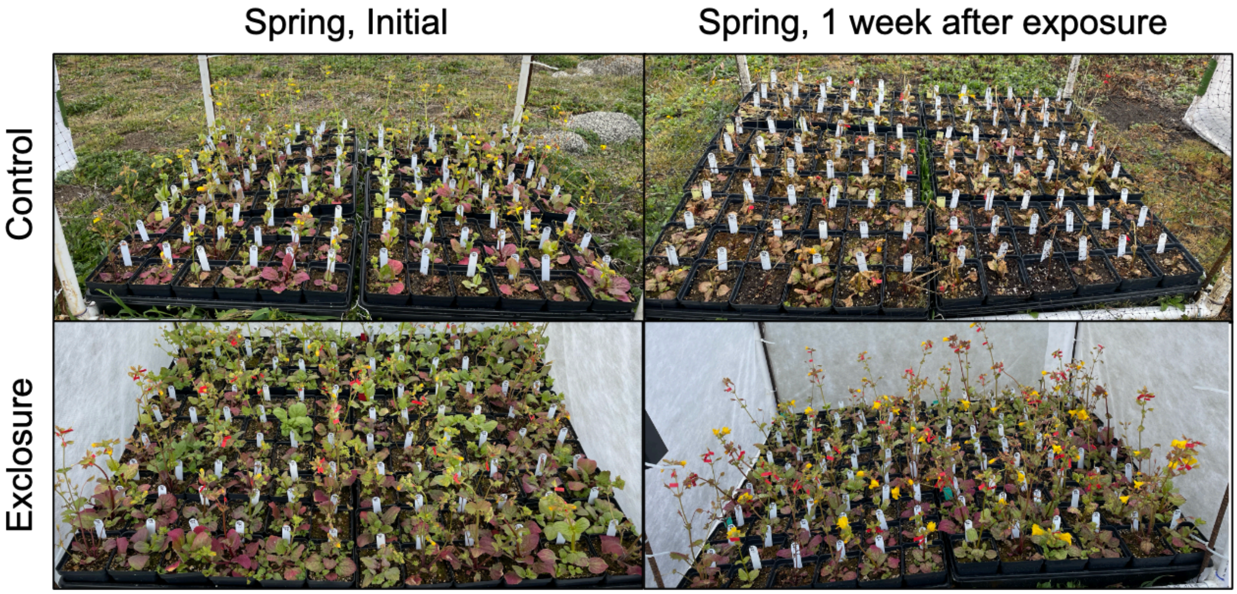


**Figure S3.** Photographs depicting a control and exclosure block in the spring cohort the day plants were first placed outside (initial) and a week later. Spring plants exposed to the ocean exhibited substantial tissue necrosis after one week, while agrofabric protected plants were largely undamaged.

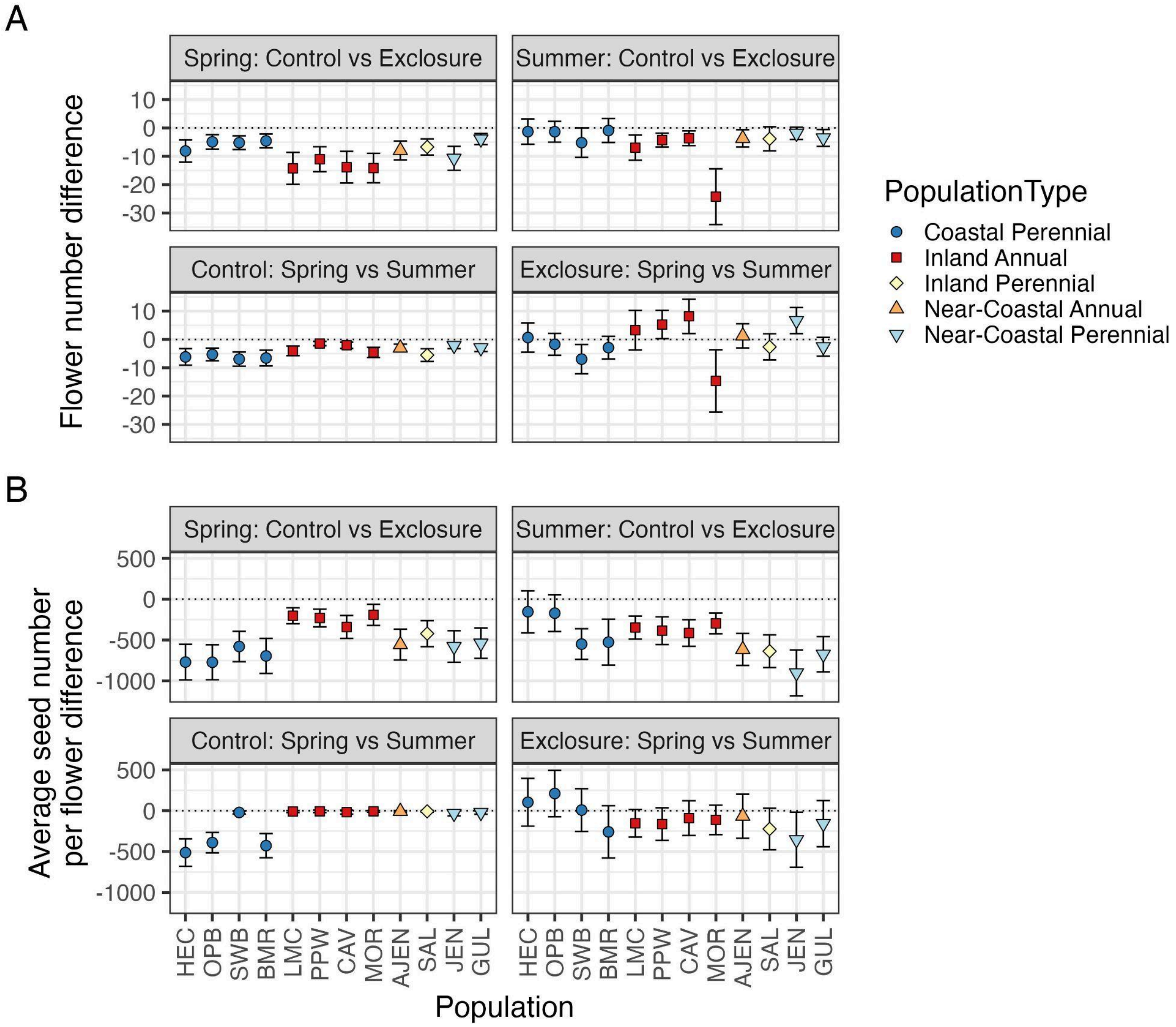


**Figure S4**. Plants produced fewer flowers (A) and seeds per flower (B) in the spring cohort compared to the summer cohort when exposed to the ocean, plants protected by agrofabric exclosures produced more flowers and seeds than those exposed to the ocean, and the benefit of this protection was greater in spring when wind speeds and salt deposition was higher. (A) Plots depict post-hoc pairwise comparison estimates from a mixed model (black outlined points; mean and 95% confidence intervals), where flower production was the response variable, population, exclosure treatment, season, all two-way interactions, and the three-way interaction were fixed predictor variables, and block nested within flat was a random effect. (B) The pairwise comparisons for seed production were from a mixed model model that included the same predictor variables as flower production except for the three-way population by season by exclosure treatment interaction. Pairwise comparisons were performed within each population for every combination of exclosure treatment and season, but only the relevant within season between exclosure treatment and within exclosure treatment between season comparisons are plotted (Full comparisons in Tables S4 and S5).

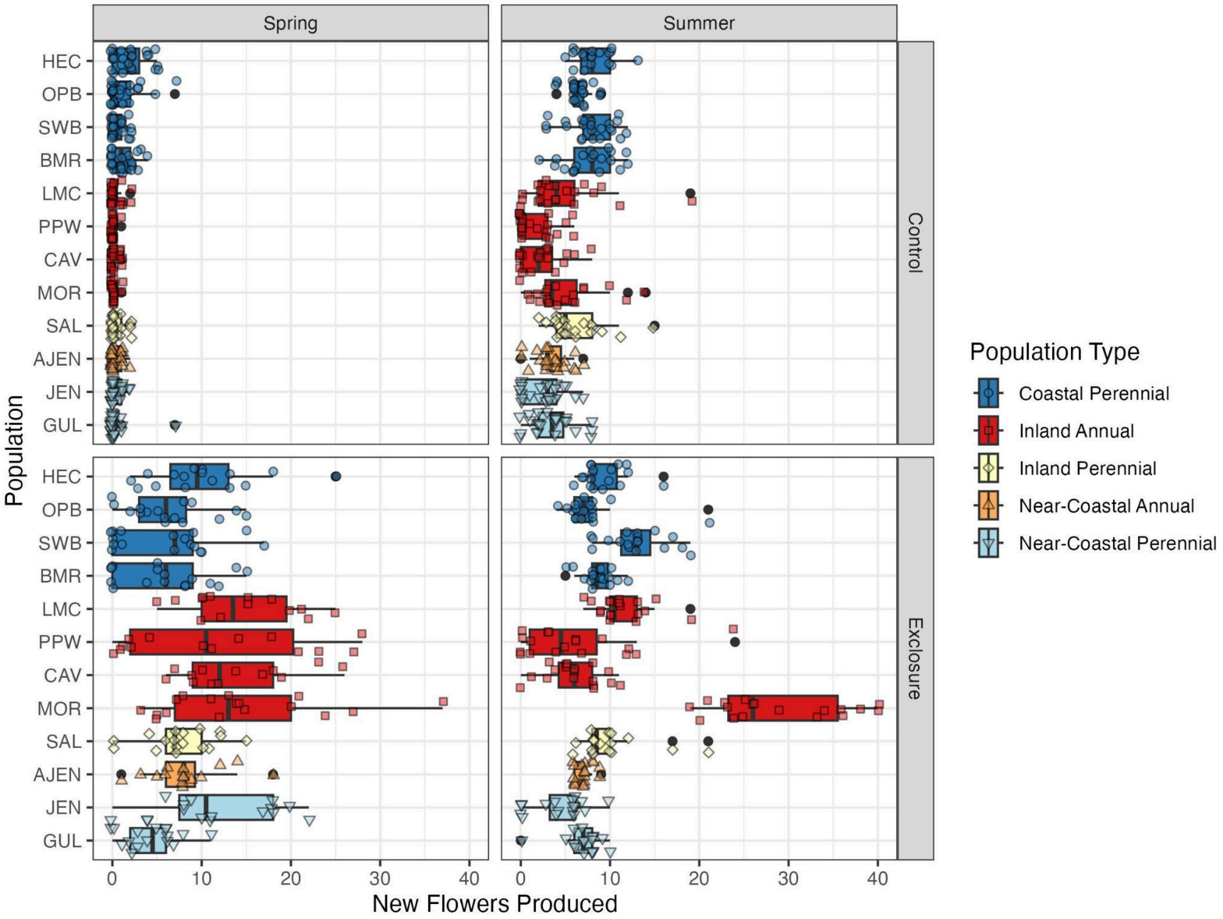


**Figure S5.** Raw values (points) and boxplots of flowers produced during the 21 day exposure period by each plant in each cohort.

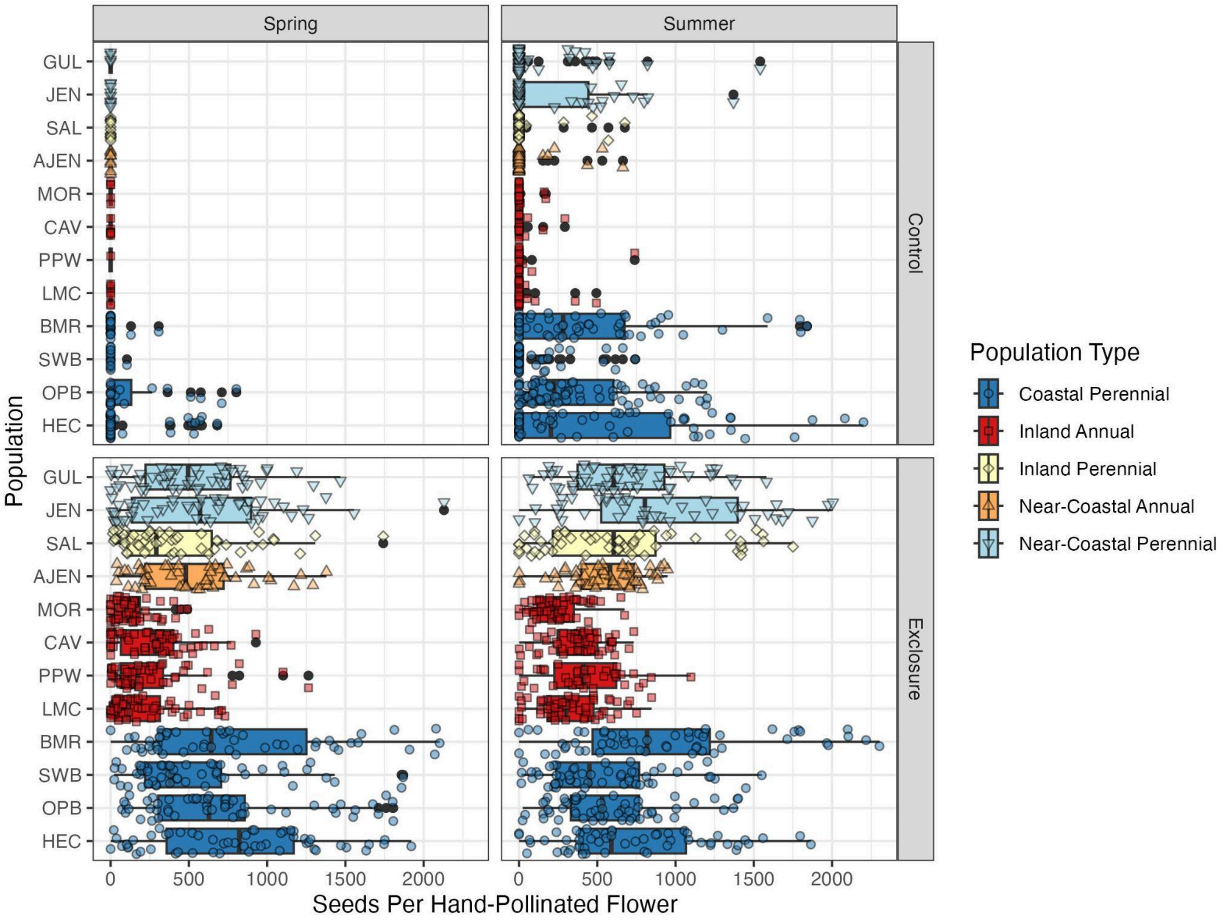


**Figure S6.** Raw values (points) and boxplots of seeds per hand-pollinated flower (up to 3 flowers per plant) produced by each plant in each cohort.

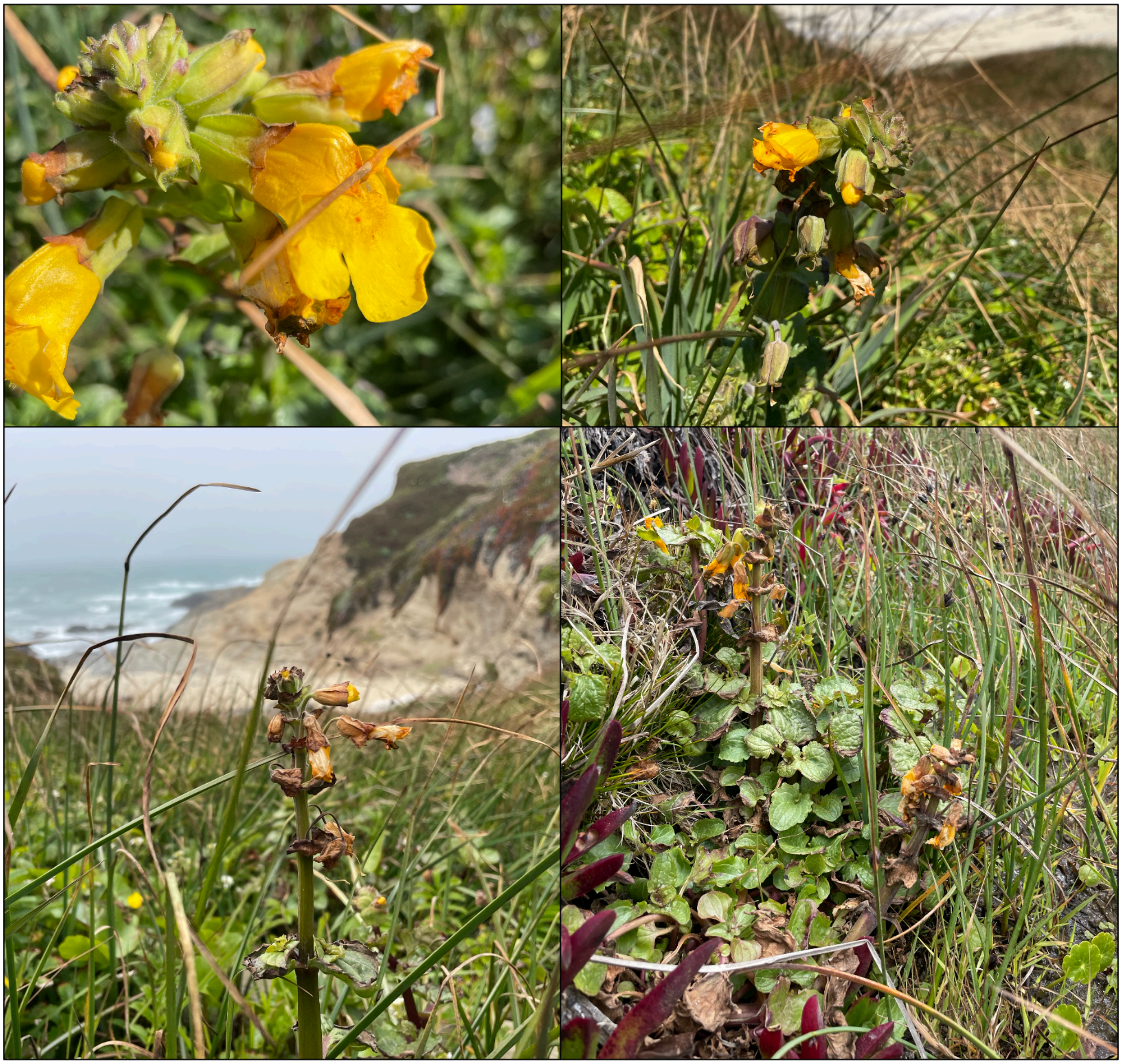

**Figure S7.** Photographs of necrosis in natural coastal perennial populations exposed to the ocean observed on Bodega Head in May 2025.

**Table S1.** Populations used in the experiments and coordinates of their original collection localities.

| Population | Population Type | Latitude | Longitude |
|---|---|---|---|
| HEC | Coastal Perennial | 44.13506 | -124.1228 |
| OPB | Coastal Perennial | 42.46401 | -124.42291 |
| SWB | Coastal Perennial | 39.03598 | -123.69046 |
| BMR | Coastal Perennial | 38.31608 | -123.06907 |
| LMC | Inland Annual | 38.86398 | -123.08391 |
| PPW | Inland Annual | 38.5755 | -122.7009 |
| CAV | Inland Annual | 38.34281 | -122.4854 |
| MOR | Inland Annual | 38.42958 | -122.94496 |
| SAL | Inland Perennial | 41.33966 | -123.38816 |
| AJEN | Near-Coastal Annual | 38.45672 | -123.11599 |
| JEN | Near-Coastal Perennial | 38.4677 | -123.12831 |
| GUL | Near-Coastal Perennial | 38.481706 | -123.13299 |

**Table S2.** Pairwise comparisons of the change in living tissue height (initial minus final height), a measure of apical necrosis. Comparisons were made from a linear mixed model consisting of population, season, exclosure treatment, all two-way interactions, and the three-way interaction as fixed factors, and flat nested within block as random factors.

| Population | contrast | estimate | SE | *df* | *z*-ratio | *p*-value |
|---|---|---|---|---|---|---|
| HEC | Spring: Control vs Exclosure | -152.347 | 10.615 | 4222 | -14.352 | 3.70E-08 |
| | Control: Spring vs Summer | -37.861 | 9.494 | 4222 | -3.988 | 0.000394544 |
| | Control Spring vs Exclosure Summer | -113.847 | 10.615 | 4222 | -10.725 | 3.70E-08 |
| | Exclosure Spring vs Control Summer | 114.486 | 10.615 | 4222 | 10.785 | 3.70E-08 |
| | Exclosure: Spring vs Summer | 38.500 | 11.628 | 4222 | 3.311 | 0.005186841 |
| | Summer: Control vs Exclosure | -75.986 | 10.615 | 4222 | -7.158 | 3.70E-08 |
| OPB | Spring: Control vs Exclosure | -74.621 | 10.615 | 4222 | -7.030 | 3.70E-08 |
| | Control: Spring vs Summer | -38.121 | 9.494 | 4222 | -4.015 | 0.000352102 |
| | Control Spring vs Exclosure Summer | -80.135 | 10.615 | 4222 | -7.549 | 3.70E-08 |
| | Exclosure Spring vs Control Summer | 36.500 | 10.615 | 4222 | 3.438 | 0.003308706 |
| | Exclosure: Spring vs Summer | -5.514 | 11.628 | 4222 | -0.474 | 0.964765518 |
| | Summer: Control vs Exclosure | -42.014 | 10.615 | 4222 | -3.958 | 0.000446205 |
| SWB | Spring: Control vs Exclosure | -186.208 | 10.615 | 4222 | -17.542 | 3.70E-08 |
| | Control: Spring vs Summer | 11.519 | 9.494 | 4222 | 1.213 | 0.61846304 |
| | Control Spring vs Exclosure Summer | -44.069 | 10.615 | 4222 | -4.152 | 0.00019722 |
| | Exclosure Spring vs Control Summer | 197.727 | 10.615 | 4222 | 18.627 | 3.70E-08 |
| | Exclosure: Spring vs Summer | 142.139 | 11.628 | 4222 | 12.224 | 3.70E-08 |
| | Summer: Control vs Exclosure | -55.588 | 10.615 | 4222 | -5.237 | 1.06E-06 |
| BMR | Spring: Control vs Exclosure | -129.435 | 10.615 | 4222 | -12.193 | 3.70E-08 |
| | Control: Spring vs Summer | -40.731 | 9.494 | 4222 | -4.290 | 0.000107502 |
| | Control Spring vs Exclosure Summer | -109.213 | 10.615 | 4222 | -10.288 | 3.70E-08 |
| | Exclosure Spring vs Control Summer | 88.704 | 10.615 | 4222 | 8.356 | 3.70E-08 |
| | Exclosure: Spring vs Summer | 20.222 | 11.628 | 4222 | 1.739 | 0.303511085 |
| | Summer: Control vs Exclosure | -68.481 | 10.615 | 4222 | -6.451 | 3.78E-08 |
| LMC | Spring: Control vs Exclosure | -51.306 | 10.615 | 4222 | -4.833 | 8.32E-06 |
| | Control: Spring vs Summer | -30.195 | 9.494 | 4222 | -3.180 | 0.008074192 |

| | | | | | | |
|---|---|---|---|---|---|---|
| | Control Spring vs Exclosure Summer | -58.792 | 10.615 | 4222 | -5.539 | 2.31E-07 |
| | Exclosure Spring vs Control Summer | 21.111 | 10.615 | 4222 | 1.989 | 0.192315624 |
| | Exclosure: Spring vs Summer | -7.486 | 11.628 | 4222 | -0.644 | 0.917731717 |
| | Summer: Control vs Exclosure | -28.597 | 10.615 | 4222 | -2.694 | 0.035684182 |
| PPW | Spring: Control vs Exclosure | -162.194 | 10.615 | 4222 | -15.280 | 3.70E-08 |
| | Control: Spring vs Summer | -22.564 | 9.494 | 4222 | -2.377 | 0.081756765 |
| | Control Spring vs Exclosure Summer | -54.499 | 10.615 | 4222 | -5.134 | 1.81E-06 |
| | Exclosure Spring vs Control Summer | 139.630 | 10.615 | 4222 | 13.154 | 3.70E-08 |
| | Exclosure: Spring vs Summer | 107.695 | 11.628 | 4222 | 9.261 | 3.70E-08 |
| | Summer: Control vs Exclosure | -31.935 | 10.615 | 4222 | -3.008 | 0.014056293 |
| CAV | Spring: Control vs Exclosure | -179.310 | 10.615 | 4222 | -16.892 | 3.70E-08 |
| | Control: Spring vs Summer | -14.157 | 9.494 | 4222 | -1.491 | 0.442942608 |
| | Control Spring vs Exclosure Summer | -75.532 | 10.615 | 4222 | -7.116 | 3.70E-08 |
| | Exclosure Spring vs Control Summer | 165.153 | 10.615 | 4222 | 15.558 | 3.70E-08 |
| | Exclosure: Spring vs Summer | 103.778 | 11.628 | 4222 | 8.925 | 3.70E-08 |
| | Summer: Control vs Exclosure | -61.375 | 10.615 | 4222 | -5.782 | 8.45E-08 |
| MOR | Spring: Control vs Exclosure | -180.296 | 10.615 | 4222 | -16.985 | 3.70E-08 |
| | Control: Spring vs Summer | -72.389 | 9.494 | 4222 | -7.624 | 3.70E-08 |
| | Control Spring vs Exclosure Summer | -173.782 | 10.615 | 4222 | -16.371 | 3.70E-08 |
| | Exclosure Spring vs Control Summer | 107.907 | 10.615 | 4222 | 10.165 | 3.70E-08 |
| | Exclosure: Spring vs Summer | 6.514 | 11.628 | 4222 | 0.560 | 0.943785097 |
| | Summer: Control vs Exclosure | -101.393 | 10.615 | 4222 | -9.552 | 3.70E-08 |
| AJEN | Spring: Control vs Exclosure | -50.454 | 10.615 | 4222 | -4.753 | 1.23E-05 |
| | Control: Spring vs Summer | -25.092 | 9.494 | 4222 | -2.643 | 0.041101817 |
| | Control Spring vs Exclosure Summer | -41.314 | 10.615 | 4222 | -3.892 | 0.000584235 |
| | Exclosure Spring vs Control Summer | 25.361 | 10.615 | 4222 | 2.389 | 0.07928762 |
| | Exclosure: Spring vs Summer | 9.139 | 11.628 | 4222 | 0.786 | 0.860866586 |
| | Summer: Control vs Exclosure | -16.222 | 10.615 | 4222 | -1.528 | 0.420612157 |
| SAL | Spring: Control vs Exclosure | -196.736 | 10.615 | 4222 | -18.534 | 3.70E-08 |
| | Control: Spring vs Summer | -9.917 | 9.494 | 4222 | -1.045 | 0.723216565 |

| | | | | | | |
|---|---|---|---|---|---|---|
| | Control Spring vs Exclosure Summer | -71.042 | 10.615 | 4222 | -6.693 | 3.72E-08 |
| | Exclosure Spring vs Control Summer | 186.819 | 10.615 | 4222 | 17.599 | 3.70E-08 |
| | Exclosure: Spring vs Summer | 125.694 | 11.628 | 4222 | 10.809 | 3.70E-08 |
| | Summer: Control vs Exclosure | -61.125 | 10.615 | 4222 | -5.758 | 9.15E-08 |
| JEN | Spring: Control vs Exclosure | -152.972 | 10.615 | 4222 | -14.411 | 3.70E-08 |
| | Control: Spring vs Summer | -138.630 | 9.494 | 4222 | -14.601 | 3.70E-08 |
| | Control Spring vs Exclosure Summer | -195.834 | 10.615 | 4222 | -18.449 | 3.70E-08 |
| | Exclosure Spring vs Control Summer | 14.343 | 10.615 | 4222 | 1.351 | 0.530290895 |
| | Exclosure: Spring vs Summer | -42.861 | 11.628 | 4222 | -3.686 | 0.001318564 |
| | Summer: Control vs Exclosure | -57.204 | 10.615 | 4222 | -5.389 | 4.84E-07 |
| GUL | Spring: Control vs Exclosure | -123.866 | 10.615 | 4222 | -11.669 | 3.70E-08 |
| | Control: Spring vs Summer | -105.055 | 9.494 | 4222 | -11.065 | 3.70E-08 |
| | Control Spring vs Exclosure Summer | -153.435 | 10.615 | 4222 | -14.454 | 3.70E-08 |
| | Exclosure Spring vs Control Summer | 18.810 | 10.615 | 4222 | 1.772 | 0.287005258 |
| | Exclosure: Spring vs Summer | -29.569 | 11.628 | 4222 | -2.543 | 0.053714451 |
| | Summer: Control vs Exclosure | -48.379 | 10.615 | 4222 | -4.558 | 3.16E-05 |

**Table S3.** Overall counts of leaf and inflorescence necrosis by the end of the spring and summer exposure periods.

| | | Leaf Necrosis | | | Inflorescence Necrosis | | |
|---|---|---|---|---|---|---|---|
| Season | Treatment | None | Partial | Complete | None | Partial | Complete |
| Spring | Control | 0 | 10 | 314 | 0 | 1 | 323 |
| | Exclosure | 190 | 20 | 6 | 196 | 6 | 14 |
| Summer | Control | 56 | 268 | 0 | 71 | 253 | 0 |
| | Exclosure | 216 | 0 | 0 | 216 | 0 | 0 |

**Table S4**. Pairwise comparisons of flowers produced over 21 days outdoors. Comparisons were made from a mixed model with a negative binomial error distribution consisting of population, season, exclosure treatment, all two-way interactions, and the three-way interaction as fixed factors, and flat nested within block as random factors.

| Population | contrast | estimate | SE | *df* | *z*-ratio | *p*-value |
|---|---|---|---|---|---|---|
| HEC | Spring: Control vs Exclosure | -8.147 | 1.532 | Inf | -5.316 | 6.33E-07 |
| | Control: Spring vs Summer | -6.176 | 1.134 | Inf | -5.446 | 3.09E-07 |
| | Control Spring vs Exclosure Summer | -7.481 | 1.390 | Inf | -5.383 | 4.37E-07 |
| | Exclosure Spring vs Control Summer | 1.971 | 1.852 | Inf | 1.064 | 0.71132193 |
| | Exclosure: Spring vs Summer | 0.666 | 2.019 | Inf | 0.330 | 0.987614849 |
| | Summer: Control vs Exclosure | -1.305 | 1.736 | Inf | -0.752 | 0.875968098 |
| OPB | Spring: Control vs Exclosure | -4.912 | 0.997 | Inf | -4.927 | 4.98E-06 |
| | Control: Spring vs Summer | -5.308 | 0.873 | Inf | -6.078 | 7.30E-09 |
| | Control Spring vs Exclosure Summer | -6.637 | 1.176 | Inf | -5.645 | 9.92E-08 |
| | Exclosure Spring vs Control Summer | -0.397 | 1.287 | Inf | -0.308 | 0.989833006 |
| | Exclosure: Spring vs Summer | -1.726 | 1.509 | Inf | -1.144 | 0.662181368 |
| | Summer: Control vs Exclosure | -1.329 | 1.430 | Inf | -0.929 | 0.789058137 |
| SWB | Spring: Control vs Exclosure | -5.201 | 0.936 | Inf | -5.558 | 1.64E-07 |
| | Control: Spring vs Summer | -6.950 | 0.974 | Inf | -7.134 | 5.86E-12 |
| | Control Spring vs Exclosure Summer | -12.151 | 1.782 | Inf | -6.819 | 5.49E-11 |
| | Exclosure Spring vs Control Summer | -1.749 | 1.333 | Inf | -1.312 | 0.555158244 |
| | Exclosure: Spring vs Summer | -6.950 | 2.001 | Inf | -3.474 | 0.002886125 |
| | Summer: Control vs Exclosure | -5.201 | 2.019 | Inf | -2.576 | 0.049027677 |
| BMR | Spring: Control vs Exclosure | -4.564 | 0.941 | Inf | -4.852 | 7.27E-06 |
| | Control: Spring vs Summer | -6.554 | 1.078 | Inf | -6.080 | 7.20E-09 |
| | Control Spring vs Exclosure Summer | -7.466 | 1.284 | Inf | -5.815 | 3.63E-08 |
| | Exclosure Spring vs Control Summer | -1.990 | 1.392 | Inf | -1.430 | 0.480391334 |
| | Exclosure: Spring vs Summer | -2.902 | 1.557 | Inf | -1.863 | 0.243977877 |
| | Summer: Control vs Exclosure | -0.911 | 1.644 | Inf | -0.554 | 0.945400374 |
| LMC | Spring: Control vs Exclosure | -14.268 | 2.193 | Inf | -6.505 | 4.66E-10 |
| | Control: Spring vs Summer | -4.039 | 0.660 | Inf | -6.117 | 5.72E-09 |

| | | | | | | |
|---|---|---|---|---|---|---|
| | Control Spring vs Exclosure Summer | -10.996 | 1.606 | Inf | -6.846 | 4.56E-11 |
| | Exclosure Spring vs Control Summer | 10.229 | 2.285 | Inf | 4.476 | 4.50E-05 |
| | Exclosure: Spring vs Summer | 3.272 | 2.714 | Inf | 1.206 | 0.623314312 |
| | Summer: Control vs Exclosure | -6.957 | 1.730 | Inf | -4.022 | 3.36E-04 |
| PPW | Spring: Control vs Exclosure | -11.038 | 1.714 | Inf | -6.439 | 7.21E-10 |
| | Control: Spring vs Summer | -1.463 | 0.288 | Inf | -5.071 | 2.36177E-06 |
| | Control Spring vs Exclosure Summer | -5.772 | 0.913 | Inf | -6.321 | 1.56E-09 |
| | Exclosure Spring vs Control Summer | 9.575 | 1.737 | Inf | 5.511 | 2.13E-07 |
| | Exclosure: Spring vs Summer | 5.266 | 1.941 | Inf | 2.712 | 0.033786907 |
| | Summer: Control vs Exclosure | -4.309 | 0.956 | Inf | -4.507 | 3.90E-05 |
| CAV | Spring: Control vs Exclosure | -13.861 | 2.175 | Inf | -6.373 | 1.11E-09 |
| | Control: Spring vs Summer | -2.058 | 0.436 | Inf | -4.722 | 1.39E-05 |
| | Control Spring vs Exclosure Summer | -5.699 | 0.929 | Inf | -6.131 | 5.23E-09 |
| | Exclosure Spring vs Control Summer | 11.802 | 2.215 | Inf | 5.329 | 5.90E-07 |
| | Exclosure: Spring vs Summer | 8.162 | 2.362 | Inf | 3.456 | 0.003080277 |
| | Summer: Control vs Exclosure | -3.640 | 1.019 | Inf | -3.573 | 0.002000462 |
| MOR | Spring: Control vs Exclosure | -14.176 | 2.020 | Inf | -7.017 | 1.36E-11 |
| | Control: Spring vs Summer | -4.573 | 0.700 | Inf | -6.532 | 3.88E-10 |
| | Control Spring vs Exclosure Summer | -28.824 | 3.776 | Inf | -7.634 | 1.61E-13 |
| | Exclosure Spring vs Control Summer | 9.603 | 2.137 | Inf | 4.494 | 4.13E-05 |
| | Exclosure: Spring vs Summer | -14.649 | 4.281 | Inf | -3.421 | 0.003485861 |
| | Summer: Control vs Exclosure | -24.252 | 3.839 | Inf | -6.316 | 1.61E-09 |
| AJEN | Spring: Control vs Exclosure | -7.956 | 1.284 | Inf | -6.195 | 3.50E-09 |
| | Control: Spring vs Summer | -3.013 | 0.528 | Inf | -5.703 | 7.07E-08 |
| | Control Spring vs Exclosure Summer | -6.697 | 1.074 | Inf | -6.238 | 2.67E-09 |
| | Exclosure Spring vs Control Summer | 4.942 | 1.379 | Inf | 3.583 | 0.00192531 |
| | Exclosure: Spring vs Summer | 1.259 | 1.666 | Inf | 0.756 | 0.874225801 |
| | Summer: Control vs Exclosure | -3.683 | 1.186 | Inf | -3.107 | 0.010217593 |
| SAL | Spring: Control vs Exclosure | -6.721 | 1.115 | Inf | -6.027 | 1.00E-08 |
| | Control: Spring vs Summer | -5.537 | 0.867 | Inf | -6.383 | 1.04E-09 |

| | | | | | | |
|---|---|---|---|---|---|---|
| | Control Spring vs Exclosure Summer | -9.366 | 1.427 | Inf | -6.564 | 3.13E-10 |
| | Exclosure Spring vs Control Summer | 1.184 | 1.398 | Inf | 0.847 | 0.831854091 |
| | Exclosure: Spring vs Summer | -2.645 | 1.800 | Inf | -1.469 | 0.456112121 |
| | Summer: Control vs Exclosure | -3.829 | 1.658 | Inf | -2.310 | 0.095746552 |
| JEN | Spring: Control vs Exclosure | -10.706 | 1.643 | Inf | -6.516 | 4.32E-10 |
| | Control: Spring vs Summer | -2.144 | 0.435 | Inf | -4.926 | 5.00E-06 |
| | Control Spring vs Exclosure Summer | -4.056 | 0.748 | Inf | -5.425 | 3.46E-07 |
| | Exclosure Spring vs Control Summer | 8.562 | 1.690 | Inf | 5.065 | 2.44E-06 |
| | Exclosure: Spring vs Summer | 6.649 | 1.796 | Inf | 3.701 | 0.001228134 |
| | Summer: Control vs Exclosure | -1.913 | 0.847 | Inf | -2.258 | 0.108002574 |
| GUL | Spring: Control vs Exclosure | -3.917 | 0.763 | Inf | -5.137 | 1.67E-06 |
| | Control: Spring vs Summer | -2.969 | 0.507 | Inf | -5.855 | 2.85E-08 |
| | Control Spring vs Exclosure Summer | -6.497 | 1.052 | Inf | -6.175 | 3.96E-09 |
| | Exclosure Spring vs Control Summer | 0.949 | 0.900 | Inf | 1.054 | 0.717368493 |
| | Exclosure: Spring vs Summer | -2.580 | 1.288 | Inf | -2.003 | 0.186911296 |
| | Summer: Control vs Exclosure | -3.528 | 1.155 | Inf | -3.054 | 0.012101607 |

**Table S5**. Pairwise comparisons of seeds produced by hand pollinated flowers outdoors. Comparisons were made from a mixed model with a zero-inflated negative binomial error distribution consisting of population, season, exclosure treatment, all two-way interactions as fixed factors, and flat nested within block as random factors.

| Population | contrast | estimate | SE | *df* | *z*-ratio | *p*-value |
|---|---|---|---|---|---|---|
| HEC | Spring: Control vs Exclosure | -769.847 | 84.949 | Inf | -9.063 | 5.2958E-14 |
| | Control: Spring vs Summer | -512.981 | 65.409 | Inf | -7.843 | 8.58E-14 |
| | Control Spring vs Exclosure Summer | -666.690 | 80.655 | Inf | -8.266 | 4.56E-14 |
| | Exclosure Spring vs Control Summer | 256.866 | 110.778 | Inf | 2.319 | 0.09375824 |
| | Exclosure: Spring vs Summer | 103.157 | 113.655 | Inf | 0.908 | 0.80078479 |
| | Summer: Control vs Exclosure | -153.709 | 100.185 | Inf | -1.534 | 0.41689778 |
| OPB | Spring: Control vs Exclosure | -772.113 | 83.518 | Inf | -9.245 | 4.341E-14 |
| | Control: Spring vs Summer | -390.766 | 48.469 | Inf | -8.062 | 5.57E-14 |
| | Control Spring vs Exclosure Summer | -561.743 | 76.304 | Inf | -7.362 | 1.12E-12 |
| | Exclosure Spring vs Control Summer | 381.347 | 99.503 | Inf | 3.833 | 0.00073196 |
| | Exclosure: Spring vs Summer | 210.370 | 110.430 | Inf | 1.905 | 0.2258433 |
| | Summer: Control vs Exclosure | -170.978 | 87.284 | Inf | -1.959 | 0.2037524 |
| SWB | Spring: Control vs Exclosure | -578.779 | 72.442 | Inf | -7.990 | 6.195E-14 |
| | Control: Spring vs Summer | -21.928 | 7.017 | Inf | -3.125 | 9.62E-03 |
| | Control Spring vs Exclosure Summer | -571.166 | 72.885 | Inf | -7.837 | 8.7375E-14 |
| | Exclosure Spring vs Control Summer | 556.851 | 72.921 | Inf | 7.636 | 1.59E-13 |
| | Exclosure: Spring vs Summer | 7.613 | 102.070 | Inf | 0.075 | 0.99985129 |
| | Summer: Control vs Exclosure | -549.238 | 72.979 | Inf | -7.526 | 3.40E-13 |
| BMR | Spring: Control vs Exclosure | -694.332 | 83.273 | Inf | -8.338 | 4.2411E-14 |
| | Control: Spring vs Summer | -427.902 | 57.812 | Inf | -7.402 | 8.3356E-13 |
| | Control Spring vs Exclosure Summer | -953.790 | 95.921 | Inf | -9.943 | 3.3973E-14 |
| | Exclosure Spring vs Control Summer | 266.430 | 104.732 | Inf | 2.544 | 0.05342768 |
| | Exclosure: Spring vs Summer | -259.458 | 124.280 | Inf | -2.088 | 0.15706887 |
| | Summer: Control vs Exclosure | -525.887 | 109.323 | Inf | -4.810 | 8.97E-06 |
| LMC | Spring: Control vs Exclosure | -202.976 | 37.722 | Inf | -5.381 | 4.4344E-07 |
| | Control: Spring vs Summer | -10.249 | 4.615 | Inf | -2.221 | 1.18E-01 |

| | | | | | | |
|---|---|---|---|---|---|---|
| | Control Spring vs Exclosure Summer | -356.911 | 54.539 | Inf | -6.544 | 3.5903E-10 |
| | Exclosure Spring vs Control Summer | 192.727 | 38.011 | Inf | 5.070 | 2.37E-06 |
| | Exclosure: Spring vs Summer | -153.935 | 65.804 | Inf | -2.339 | 0.08928055 |
| | Summer: Control vs Exclosure | -346.662 | 54.640 | Inf | -6.344 | 1.34E-09 |
| PPW | Spring: Control vs Exclosure | -229.247 | 42.063 | Inf | -5.450 | 3.01E-07 |
| | Control: Spring vs Summer | -7.934 | 5.628 | Inf | -1.410 | 4.93E-01 |
| | Control Spring vs Exclosure Summer | -393.488 | 65.780 | Inf | -5.982 | 1.32E-08 |
| | Exclosure Spring vs Control Summer | 221.312 | 42.449 | Inf | 5.214 | 1.11E-06 |
| | Exclosure: Spring vs Summer | -164.242 | 77.635 | Inf | -2.116 | 0.14808177 |
| | Summer: Control vs Exclosure | -385.554 | 65.960 | Inf | -5.845 | 3.03E-08 |
| CAV | Spring: Control vs Exclosure | -340.107 | 54.670 | Inf | -6.221 | 2.96E-09 |
| | Control: Spring vs Summer | -18.136 | 9.115 | Inf | -1.990 | 1.92E-01 |
| | Control Spring vs Exclosure Summer | -431.523 | 63.040 | Inf | -6.845 | 4.58E-11 |
| | Exclosure Spring vs Control Summer | 321.971 | 55.575 | Inf | 5.793 | 4.13E-08 |
| | Exclosure: Spring vs Summer | -91.417 | 82.876 | Inf | -1.103 | 0.68764994 |
| | Summer: Control vs Exclosure | -413.387 | 63.489 | Inf | -6.511 | 4.47E-10 |
| MOR | Spring: Control vs Exclosure | -191.599 | 50.124 | Inf | -3.823 | 0.00076186 |
| | Control: Spring vs Summer | -7.974 | 4.014 | Inf | -1.987 | 1.93E-01 |
| | Control Spring vs Exclosure Summer | -304.501 | 49.231 | Inf | -6.185 | 3.72E-09 |
| | Exclosure Spring vs Control Summer | 183.625 | 50.365 | Inf | 3.646 | 1.52E-03 |
| | Exclosure: Spring vs Summer | -112.901 | 70.462 | Inf | -1.602 | 0.3772626 |
| | Summer: Control vs Exclosure | -296.527 | 49.309 | Inf | -6.014 | 1.09E-08 |
| AJEN | Spring: Control vs Exclosure | -555.298 | 73.290 | Inf | -7.577 | 2.3803E-13 |
| | Control: Spring vs Summer | -7.486 | 3.766 | Inf | -1.988 | 1.93E-01 |
| | Control Spring vs Exclosure Summer | -622.570 | 76.153 | Inf | -8.175 | 4.9072E-14 |
| | Exclosure Spring vs Control Summer | 547.811 | 73.417 | Inf | 7.462 | 5.39E-13 |
| | Exclosure: Spring vs Summer | -67.272 | 105.167 | Inf | -0.640 | 0.91915904 |
| | Summer: Control vs Exclosure | -615.083 | 76.174 | Inf | -8.075 | 5.50E-14 |
| SAL | Spring: Control vs Exclosure | -421.320 | 62.146 | Inf | -6.780 | 7.2366E-11 |
| | Control: Spring vs Summer | -7.996 | 4.022 | Inf | -1.988 | 1.92E-01 |

| | | | | | | |
|---|---|---|---|---|---|---|
| | Control Spring vs Exclosure Summer | -644.898 | 77.710 | Inf | -8.299 | 4.4298E-14 |
| | Exclosure Spring vs Control Summer | 413.324 | 62.298 | Inf | 6.635 | 1.95E-10 |
| | Exclosure: Spring vs Summer | -223.577 | 98.980 | Inf | -2.259 | 0.10783097 |
| | Summer: Control vs Exclosure | -636.902 | 77.746 | Inf | -8.192 | 4.82E-14 |
| JEN | Spring: Control vs Exclosure | -579.641 | 75.149 | Inf | -7.713 | 9.7478E-14 |
| | Control: Spring vs Summer | -33.237 | 11.817 | Inf | -2.813 | 2.53E-02 |
| | Control Spring vs Exclosure Summer | -935.457 | 108.450 | Inf | -8.626 | 3.4861E-14 |
| | Exclosure Spring vs Control Summer | 546.404 | 76.253 | Inf | 7.166 | 4.67E-12 |
| | Exclosure: Spring vs Summer | -355.817 | 131.302 | Inf | -2.710 | 0.03401799 |
| | Summer: Control vs Exclosure | -902.221 | 108.869 | Inf | -8.287 | 4.42E-14 |
| GUL | Spring: Control vs Exclosure | -538.031 | 71.945 | Inf | -7.478 | 4.7728E-13 |
| | Control: Spring vs Summer | -22.487 | 8.009 | Inf | -2.808 | 2.57E-02 |
| | Control Spring vs Exclosure Summer | -696.362 | 83.628 | Inf | -8.327 | 4.2966E-14 |
| | Exclosure Spring vs Control Summer | 515.544 | 72.469 | Inf | 7.114 | 6.79E-12 |
| | Exclosure: Spring vs Summer | -158.332 | 109.794 | Inf | -1.442 | 0.47298147 |
| | Summer: Control vs Exclosure | -673.875 | 83.855 | Inf | -8.036 | 5.76E-14 |
